# OVarFlow: a resource optimized GATK 4 based Open source Variant calling workFlow

**DOI:** 10.1101/2021.05.12.443585

**Authors:** Jochen Bathke, Gesine Lühken

**Affiliations:** Institute of Animal Breeding and Genetics, Justus Liebig University Gießen, Ludwigstraße 21, 35390 Gießen, Germany

**Keywords:** Variant calling, SNP, indel, GATK, next generation sequencing, reproducibility, data parallelization, benchmarking, Java

## Abstract

**Background:** Next generation sequencing technologies are opening new doors to researchers. One application is the direct discovery of sequence variants that are causative for a phenotypic trait or a disease. The detection of an organisms alterations from a reference genome is know as variant calling, a computational task involving a complex chain of software applications. One key player in the field is the Genome Analysis Toolkit (GATK). The GATK Best Practices are commonly referred recipe for variant calling on human sequencing data. Still the fact the Best Practices are highly specialized on human sequencing data and are permanently evolving is often ignored. Reproducibility is thereby aggravated, leading to continuous reinvention of pretended GATK Best Practice workflows.

**Results:** Here we present an automatized variant calling workflow, for the detection of SNPs and indels, that is broadly applicable for model as well as non-model diploid organisms. It is derived from the GATK Best Practice workflow for “Germline short variant discovery”, without being focused on human sequencing data. The workflow has been highly optimized to achieve parallelized data evaluation and also maximize performance of individual applications to shorten overall analysis time. Optimized Java garbage collection and heap size settings for the GATK applications SortSam, MarkDuplicates, HaplotypeCaller and GatherVcfs were determined by thorough benchmarking. In doing so, runtimes of an example data evaluation could be reduced from 67 h to less than 35 h.

**Conclusions:** The demand for standardized variant calling workflows is proportionally growing with the dropping costs of next generation sequencing methods. Our workflow perfectly fits into this niche, offering automatization, reproducibility and documentation of the variant calling process. Moreover resource usage is lowered to a minimum. Thereby variant calling projects should become more standardized, reducing the barrier further for smaller institutions or groups.

## Background

Evolution and therefore the diversity of life is based upon genetic variability. This may be due to small changes in an organism’s nucleotide sequence, larger rearrangements, recombination of homologous chromosomes during chromosomal crossover or chromosome reshuffling during meiosis and sexual reproduction [1, 2, 3]. Thereby new phenotypic traits may be realized within an individual. The identification of the genetic basis of such phenotypic traits is a pivotal point of genetic studies. Especially the advent of next generation sequencing (NGS) technologies have started a new era of sequencing based genetic variant identification [4, 5]. Continuous improvements in second and third generation sequencing technologies promoted permanently declining sequencing cost, even outpacing the technological advances in the semiconductor industry described by Moore’s law [6]. This paved the way for new applications of NGS methods and a broader application thereof. In this regard NGS is even challenging SNP genotyping arrays for various genome analysis applications [7]. Whole genome (WGS) as well as whole exome sequencing (WES) are now commonly used for variant discovery with a trend to go for WGS [8] eventhough WES is still in the game [9]. Also genome-wide association studies (GWAS), which have been a domain of SNP genotyping arrays [10], are increasingly conducted using WGS [11, 12].

Generating the sequencing data is only the first step in any related research project. Major steps of the subsequent analysis include read mapping, variant calling, variant filtration and functional annotation of the reads [13, 14]. Over the last decade a plethora of variant callers have been developed [15]. The Genome Analysis Tool Kit (GATK) is among the most broadly used applications [16] and GATK Best Practices workflows are considered a kind of gold standard in the field [17, 18, 19]. The GATK includes hundreds of different tools and the GATK Best Practices are supposed to guide users through their application [17, 13]. Therefore it has become customary to simply cite the GATK Best Practices in method sections of publications while supplying a link to the GATK website [20, 21, 22]. The problems resulting from this routine are twofold. Firstly, the GATK Best Practices are a dynamic document, were command lines, arguments and tool choices can become obsolete. Years from the initial publication it might become elusive what the GATK Best Practices were by that time. Secondly, simply citing the GATK Best Practices is used as a shortcut for method sections. This has also been noticed by the developers of GATK, therefore stating on their website: “Lots of workflows that people call GATK Best Practices diverge significantly from our recommendations [23].” Reproducibility of the actual data evaluation is thereby obscured, which relates to the fact that the “Best Practices workflows […] are designed specifically for human genome research [23]” even though the GATK has successfully been used to analyze various species [20, 22, 24, 25].

A lot of the scientific work is unnecessarily time consuming due to the lack of reproducibility because of unclear methods [26]. Reproducibility is of an equal concern in the field of computational biology as for the laboratory work [27]. Especially complex workflows like variant calling and software stacks like the GATK constitute a challenge to small scientific groups and newcomers to the field.

The rapidly growing adaptation of NGS base variant calling, the broad use of the GATK and the need for reproducible data analysis highlight a demand for broadly applicable, well documented and readily usable GATK-based variant calling workflows. We therefore developed OVarFlow, an open source, highly optimized, full-fledged variant calling workflow, that generates functionally annotated variants. The workflow is highly automated and reproducible, requiring minimal user interaction. Only short read sequencing data (e.g. Illumina), a reference genome as well as annotation have to be provided.

## Implementation

Analysis of sequencing data is highly dependent on software tools, where usability, installability and archival stability is one key aspect for the usefulness of the software tools [28]. A systematic analysis showed, that a large proportion of published tools cannot readily be installed due to problems in the implementation [28].

To circumvent such nuisance OVarFlow comes with a comprehensive documentation and builds upon established technologies. This includes Conda [29] and Bioconda [30] as an environment and package manager, thereby allowing for easy installation of required software components. To orchestrate the various data processing steps, Snakemake is utilized as a workflow management engine [31]. Alternatively all required software components come bundled via container virtualization as Docker [32] or Singularity [33] containers, if manual intervention during setup is not desirable. Data processing itself is primarily relying on the GATK [16].

The complexity of computational methods is ever-growing. We are therefore convinced that a thorough documentation is vital to the usability of any software. For this reason a comprehensive documentation is an integral part of OVarFlow. It is available at “Read the Docs”, explaining in detail setup, usage, resource requirements and the single steps of the workflow.

### Overview of the workflow

A flowchart of the actual variant calling workflow is depicted in Fig. 1. The entire workflow consists of two separate phases. The basic variant calling workflow can already stand for itself, while an optional second extended workflow allows for further refinement of the called variants via base quality score recalibration (BQSR).

**Fig. 1:**
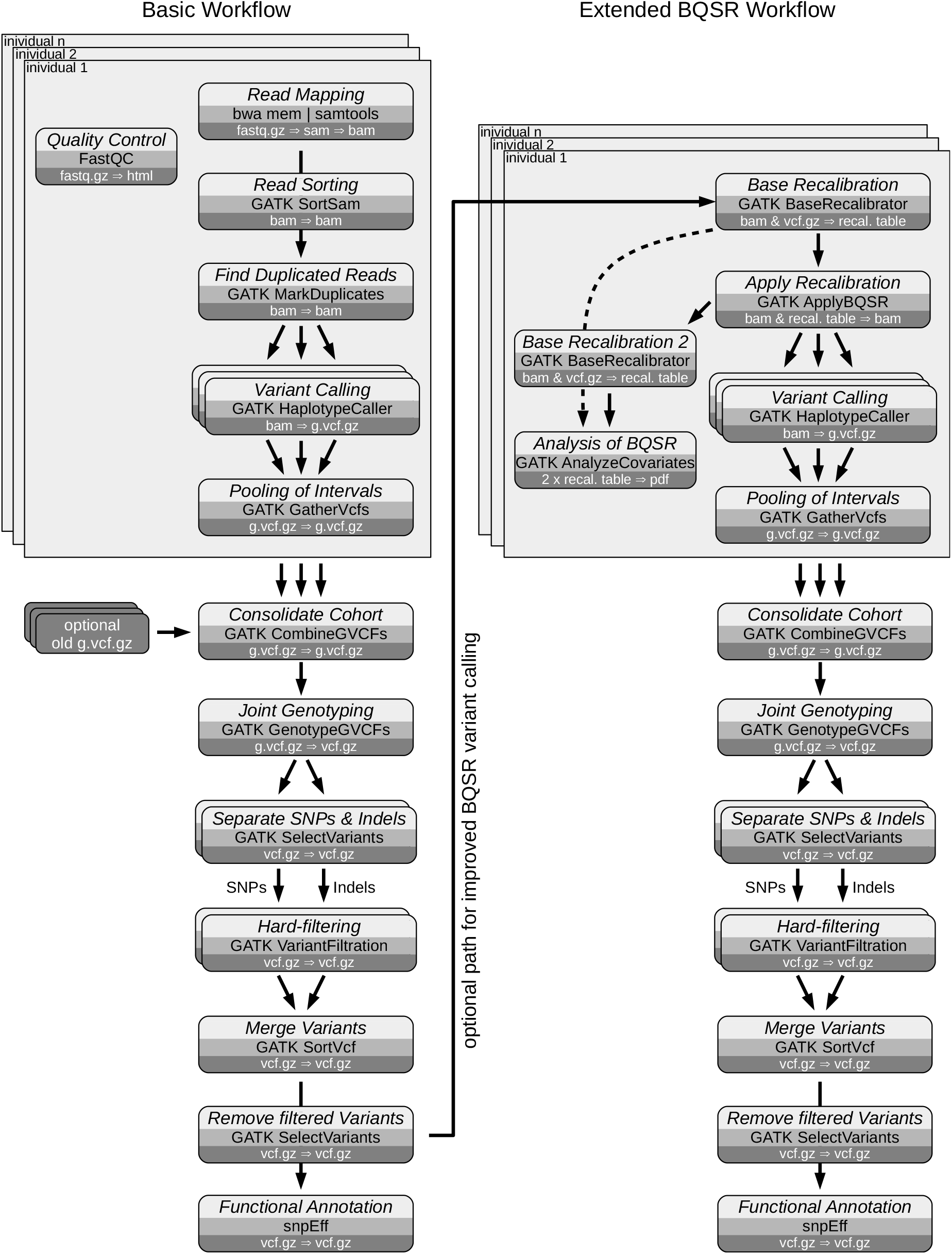
Flowchart of the variant calling workflow. The variant calling workflow consists of two separate phases. A basic workflow already generates a set of usable, functionally annotated variants (SNPs and indels). A second, optional workflow, uses the previously called variants to perform Base Quality Score Recalibration (BQSR) to improve initial base calls of the fastq files. Processing of each individuals fastq files can be performed in parallel. Also various steps of the workflow can be parallelized, e.g. base calling on genomic intervals by the GATK HaplotypeCaller, as indicated by overlapping boxes. Each box includes a description of the step (light gray), the name of the used application (medium gray) and the primary input and output data formats (dark gray).

The basic workflow requires a reference genome in fasta format (.fa.gz, optionally compressed) and reference annotation in general feature format (.gff.gz, also compressed). Finally, second generation sequencing (SGS) data have to be supplied in fastq format (.fastq.gz). Providing those data a full set of functionally annotated variants is generated. Each step of the analysis is depicted in a rounded box, naming the analysis performed, the used application and the most important input and output data. The sequencing data of several individuals can be processed in parallel and are then consolidated into a single cohort, resulting in a genomics variant call format (VCF) file (g.vcf.gz). Hard-filtering is then employed to minimize the amount of false positive variants. Finally SnpEff performs functional annotation of the called variants [34].

A second workflow can be executed in succession to the first. Thereby the called variants can further be refined through BQSR, as initially shown by DePristo *et al*. [17]. Basically BQSR allows to improve the quality scores generated by the sequencer. Still depending on the given data set, improvements achieved through BQSR may be marginal, while introducing high computational cost [35]. Therefore BQSR was included as an optional step, that might be applied to low-coverage data or badly calibrated base quality [35].

### Optimized parallelization of the workflow

Variant calling is a computationally demanding task. To speed up the analysis, parallelization was enabled wherever possible. Firstly fastq files of various individuals can be processed in parallel up to the point where variants are consolidated into a single genomics VCF file (.g.vcf.gz). Also filtering of SNPs and INDELs is performed in parallel. The two most demanding steps of the workflow are mapping via the maximal exact matches (mem) algorithm of the Burrows-Wheeler Aligner (bwa) [36] and variant calling via the GATK HaplotypeCaller. Mapping via bwa can easily be parallelized and uses a default of six threads in OVarFlow (configurable). The HaplotypeCaller on the other hand used to be parallelizable through command line switches (`-nct` and `-nc`), but those options where abandoned with GATK 4. As an alternative the GATK team introduced Spark for multithreading [37]. At the time of writing, HaplotypeCallerSpark (version 4.2.0 and below) “is still under development and should not be used for production work”, as stated by the developers [38]. OVarFlow reintroduces parallelization to the HaplotypeCaller by splitting the reference genome into several intervals, on which variant calling is performed in parallel. This approach is also called “scatter gather”. The amount of intervals processed in parallel is configurable, but defaults to four. A python script, that is part of the workflow, splits the reference genome between individual fasta sequences into intervals. Individual contigs are not split. Furthermore resource usage of the HaplotypeCaller with different numbers of native pairHMM threads was assessed (see results). While four native pairHMM threads gave the best runtime for the individual HaplotypeCaller process, more processes could be run in parallel when only a single native pairHMM thread was enabled. This was made configurable in the workflow, while defaulting to four native pairHMM threads. Finally the entire analysis scales with the given hardware. Here Snakemake helps to achieve an optimized resource utilization and parallelization from high end desktop over server to cluster usage [31].

### Pitfalls when working with the GATK

The GATK is a very complex framework, with hundreds of applications related to variant calling. Many of those show some very intricate and subtile usability issues, that are challenging to novel but also to expert users. One of those pitfalls involves hard filtering, which is often employed to remove false positive variants [39, 13]. Often hard filtering is applied by chaining several filtering expressions through the logical or operator “`||`” [25]. This might be due to a publication by the authors of GATK in 2013, where this was suggested [13]. This procedure has changed. As of today the GATK teams states, “it is strongly recommended that such expressions be provided as individual arguments … [to] ensures that all of the individual expression are applied to each variant” [40]. Therefore OVarFlow applies a separate filter to each filtration threshold (for SNPs: QD < 2.0, QUAL <30.0, SOR > 3.0, FS > 60.0, MQ < 40.0, MQRankSum < −12.5, ReadPosRankSum < −8.0; for INDELs: QD < 2.0, QUAL < 30.0, FS > 200.0, ReadPosRankSum < - 20.0).

Another potential pitfall is the CPU instruction set. Performance of the GATK 4 was optimized in an collaboration between Intel and the Broad Institute. This includes usage of Advanced Vector Extensions (AVX) for the HaplotypeCaller [41, 42]. The lack of AVX will drastically slow down the HaplotypeCaller (personal estimates are approx. 5-fold longer runtimes). OVarFlow therefore verifies the availability of AVX before executing the workflow, and informs the user about the absence of AVX. Fortunately AVX evolved to be common in newer CPU generations.

The GATK is written in the Java programming language, which is inherently linked to the Java Virtual Machine (JVM). Performance of the JVM can be optimized through several hundreds of settings (see: `java -XX:+PrintFlagsFinal` and `java -X`). Especially the Java heap size and number of garbage collection threads exert a major influence on the performance of various GATK applications (see results section). To achieve an optimized resource utilization and run time, those two parameters were optimized for the most important tools running in parallel and incorporated into the workflow.

Moreover several smaller inconveniences are taken care of by the workflow. This includes the storage of intermediate data under the `/tmp` direcotry. Depending on the partition scheme and size, this can be a source of major trouble. Therefore temporary data are stored under `GATK_tmp_dir` within the project directory. Depending on the input data the GATK application MarkDuplicates opens a plethora of files, which is even more problematic as several instances of MarkDuplicates can run in parallel. To prevent problems resulting from the number of maximum allowed open file descriptos (see `ulimit -Sn` or `-Hn`), each instance of MarkDuplicates was limited to use 300 file handles. Finally the workflow relieves the user of having to fiddle around with many intermediate files including various indices (`.bai`, `.fai`, .`dict`, `bwa` index), conversion of the gzip to BGZ format [43] or creation of a SnpEff database.

### The configuration files

The user is freed of as much manual intervention as possible, all while scaling over various infrastructure sizes. This streamlining of the data evaluation is also achieved through “convention over configuration”. However, configurability is implemented through two configuration files, of which one is entirely optional.

The first configuration file (`samples_and_read_groups.csv`) describes the input data. This file specifies the more “biological” data, like the reference genome and annotation as well as the sequenced samples. It is in colon separated values (csv) format, which allows for easy editing also in common spread sheet applications. Furthermore read group data have to be specified in this file. Additionally previously called variants can be included in the analysis. Also a minimum sequence length can be specified. Many genomes contain a high number of small contigs. Those can be excluded form the analysis by setting the desired cut-off value.

A second, optional configuration files (`config.yaml`) is more about the “technical” side of the workflow. If present, this file is automatically picked up by the Snakemake workflow management system. Here Java heap size and the number of garbage collection threads can be adjusted if required. Parallelization can also be configured for the number of bwa threads, intervals of the genome that can be processed in parallel through the HaplotypeCaller and the number of native pairHMM threads used by the HaplotypeCaller. Finally storage of temporary data and the maximum number of file handles used by MarkDuplicates can be configured. For the BQSR workflow a separate configuration file is available (`config.yaml`), that serves identical purposes.

## Results

The primary goal of our workflow is that it shall be of practical use in variant calling projects. To show its validity we reproduced the variant identification of a previous study. Another matter of concern was the resource efficiency of the variant identification. The potential to improve performance of GATK based variant calling was previously shown [42]. Here we are building upon those findings and further extend them into a streamlined variant calling workflow. For that reason we investigated how to optimize resource utilization on the level of single GATK applications and also on the level of the entire workflow.

### Proof of concept

To confirm the feasibility of our workflow, we reproduced the identification of a variant responsible for recessive autosomal dwarfism (adw) in chicken, as originally conducted by Wu *et al*. [44]. The adw variant was know to be located on chromosome 1, within 52-56 Mb. The identification of the variant was performed using WGS data of a single *adw/adw* individual and 261 non-affected White Leghorns as control. Within the candidate region a total of 146,070 variants could be identified, which were further reduced by various filtering steps to 11 potential candidates of which only one was categorized as a high impact variant (stop gained).

To repeat this analysis we obtained the sequencing data of the *adw/adw* individual from the European Nucleotide Archive (ENA, www.ebi.ac.uk/ena), together with raw reads of another 25 normal White Leghorns. The fastq sequencing data of those 26 chicken served as input data for the here presented variant calling workflow. To retain comparable genome coordinates the same reference genome (Gallus_gallus-5.0) was utilized as in the previous study. Following steps were similar to the data evaluation procedure conducted by Wu *et al*. Briefly, all variants within the candidate region of 52-56 Mb on chromosome 1 were extracted. All variants that were not homozygous and also not exclusive to the dwarf chicken were removed from the dataset. This resulted in a total of 1,090 variants. Those variants, that were categorized by SnpEff as moderate or high impact variants were finally selected, giving a total of 6 potential candidate variants (see Tab. 1). Most of those variants posses multiple annotations and exceed different impacts on the respective annotation. The identified candidate variant set is not identical to the 11 candidates that were found by Wu *et al*., which is not surprising giving the fact that different White Leghorns were used as a reference dataset as compared to the original study. Variants identified in both studies as potential candidates are marked with a small hook, those exclusive to our analysis with a cross. However all of those 11 variants selected by Wu *et al*. could be identified in our dataset prior to the filtering step (only the SNP at position 52,195,787 was detected as an deletion at position 52,195,786). More importantly, the causative variant (position 53,688,583) that was finally identified by Wu *et al*. was among the 6 candidates identified by our analysis.

**Tab 1:**
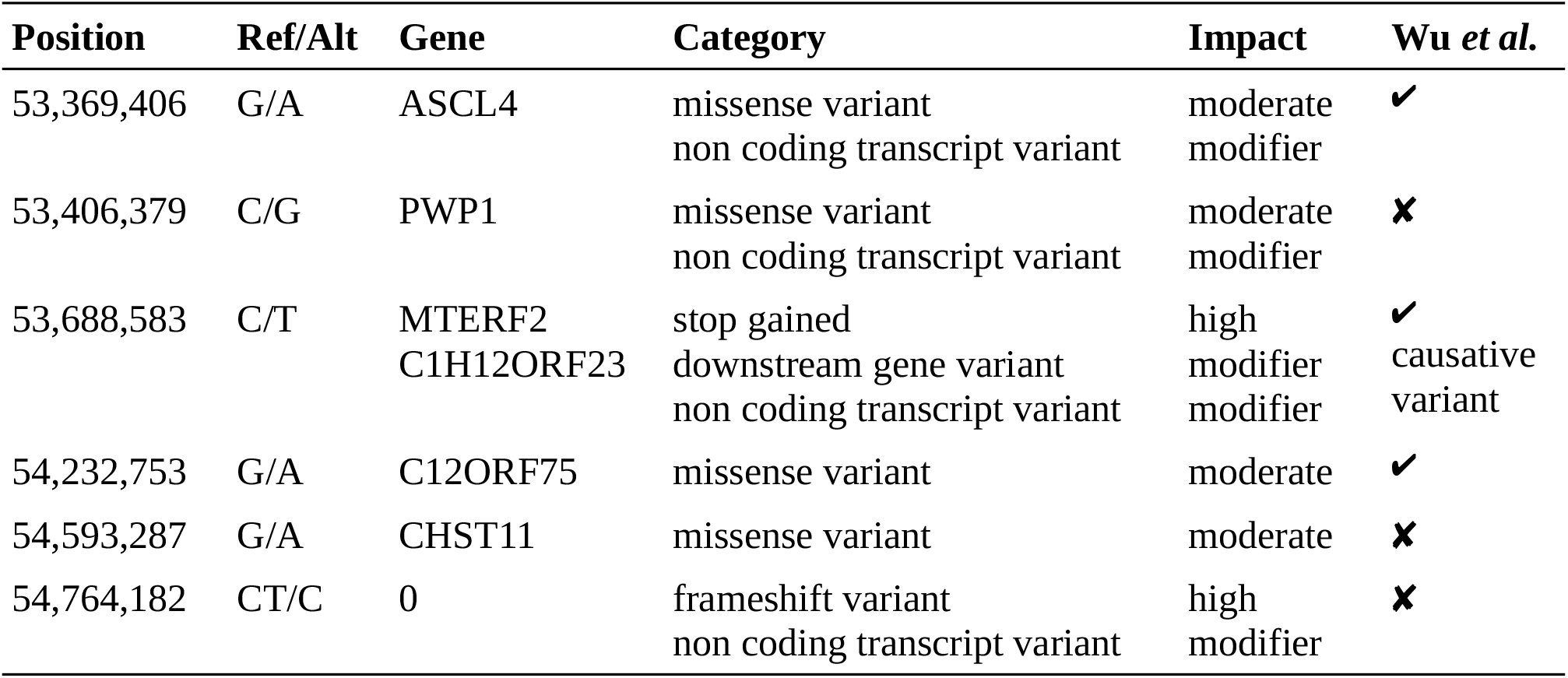
Variants exclusively associated with autosomal dwarfism in chicken.

Taken together these results confirm that our variant calling workflow is capable of detecting small variants (SNPs and indels) in whole genome sequencing data, that can be utilized in further analysis steps to identify potential causative variants.

### Optimization of individual GATK applications

The GATK is written in the Java programming language, whose bytecode is executed by the JVM. One problem with the JVM is, that its resource utilization does not always scale positively with the size of given hardware resources. This is especially problematic, as variant calling is resource hungry and demands large computational resources.

Two aspects of the JVM are automatically adjusted to the available hardware, being the number of garbage collection (GC) threads and the heap size (see Tab. 2). Both of these aspects relate to memory management through the JVM (here version 8). They were measured on actually available hardware. While the number GC threads grows continuously with the number of given CPU cores the heap size maxes out at 26.67 Gb. In many cases those numbers are considerably larger than what is needed for the respective application to run efficiently, as will be shown in the following measurements. This also means that GATK applications will posses inconsistent behavior, depending on the given hardware.

**Tab 2:**
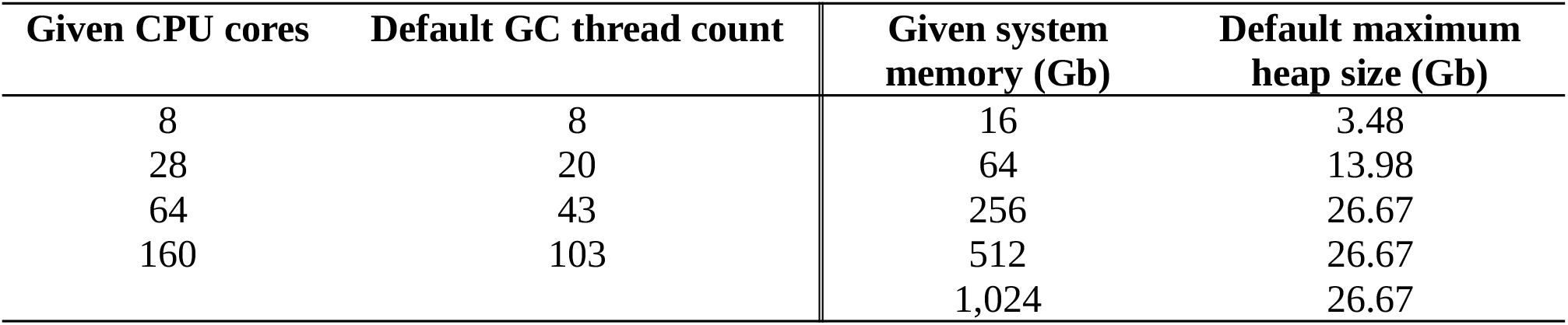
Default resource usage of the Java Virtual Machine version 8.

First we assessed the impact of GC thread count of several GATK applications (Fig. 2). We selected those applications for a deeper analysis, that are run in parallel. This part of the workflow will cause the highest load and will benefit the most from optimization. Three aspects of each application execution where analyzed, being overall execution speed (wall time), total load caused (system time) and memory usage (resident set size; RSS). Wall time of GATK SortSam, HaplotypeCaller and GatherVcfs is barely influenced by the number of GC threads. MarkDuplicates shows a linear relation, resulting in longer runtimes with higher thread counts and a sweet spot at two GC threads. Total CPU usage for SortSam and MarkDuplicates are rising with GC numbers. SortSam requires approx 50 % more CPU usage between low thread counts and 20 GC threads. The impact on MarkDuplicates is even more pronounced, resulting in an approx. 5-fold higher resource useage between 1 and 20 GC threads. No equally clear tendency is seen for HaplotypeCaller or GatherVcfs. Besides statistic variation, differences in memory consumption are not as pronounced. The GATK HaplotypeCaller might benefit from two GC threads.

**Fig. 2:**
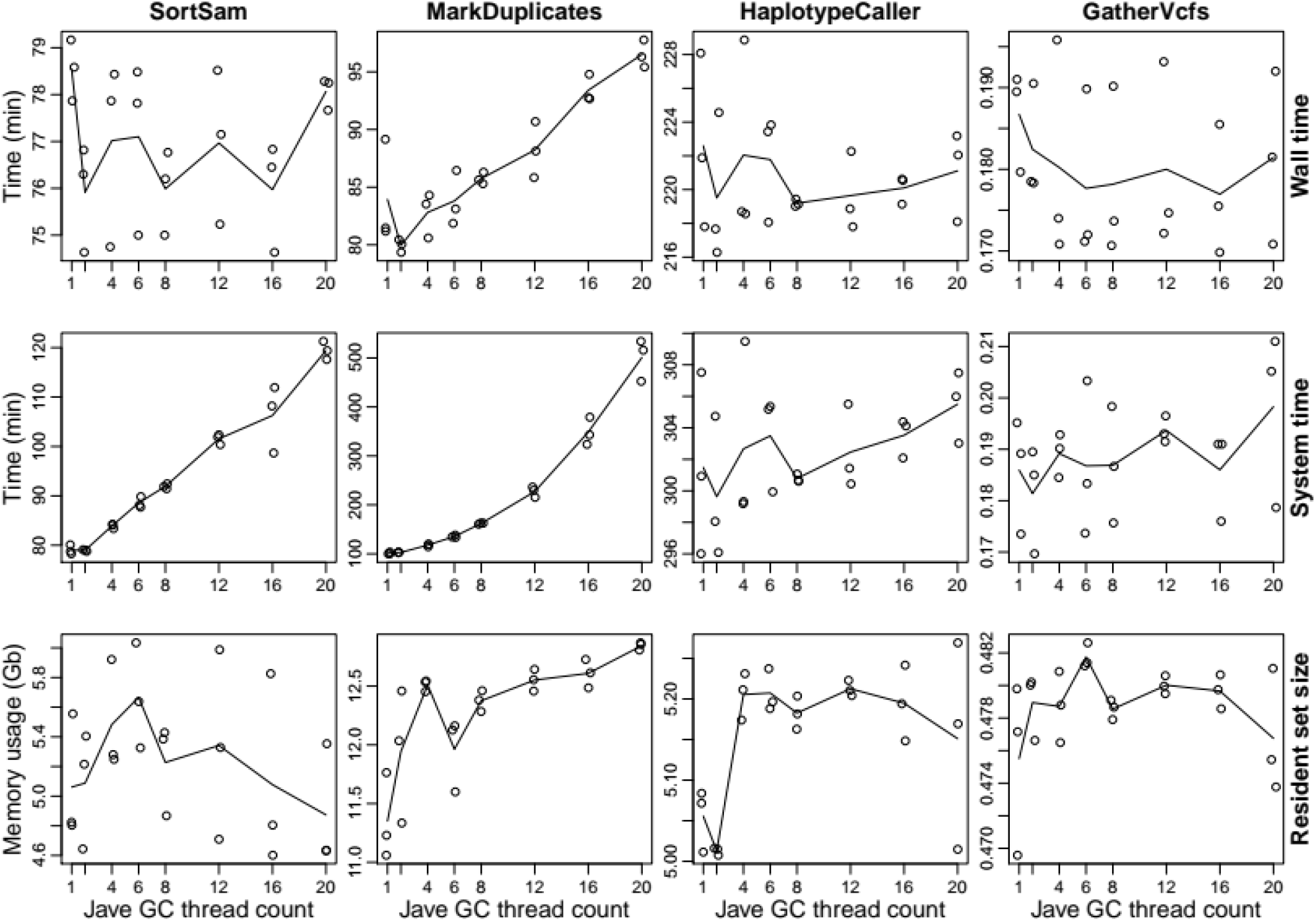
Resource usage of different GATK applications depending on Java garbage collection threads. The performance of some GATK applications is severely influenced by the number of employed Java garbage collection (GC) threads. Here the Java 8 default Paralle Garbage Collector was used. Resource usage concerning wall time, system time and resident set size (memory usage) was analyzed (see rows) for the four tools SortSam, MarkDuplicates, HaplotypeCaller and GatherVcfs (see columns) (GATK version 4.1.9). Triplicated measurements for each of eight different numbers of GC thread counts (1, 2, 4, 6, 8, 12, 16 and 20) were recorded and resulting mean values plotted in lines. Lower measured values are preferable as they reflect a lower resource usage of the respective application. The different scales of the ordinates of each plot have to be considered, as they vary greatly between the individual plots.

From those findings it can be concluded, that memory footprint is only marginally influenced by GC thread numbers. CPU load on the other side is significantly influenced for SortSam and MarkDuplicates, with a sweet spot of two GC threads, each.

Next we investigated the effects of different JVM heap sizes on the performance of the same applications, observing identical parameters as before (Fig. 3). The picture is clearly distinct. Effects on wall and system time are identical for each individual application. Especially CPU usage of SortSam benefits form larger heap sizes. Here 12 Gb are the sweet spot for the given data set. MarkDuplicates seems to benefit slightly form smaller heap sizes, but due to statistical nuisance no distinct number can be named. Memory footprint is severely affected by the maximum allowed heap size for SortSam, MarkDuplicates and HaplotypeCaller. Especially MarkDuplicates scales linearly with the maximum allowed heap size, making use of all the memory dedicated as heap space (grey line). This is especially noteworthy, as CPU usage and run time of the applications do not benefit from larger heap sizes. With SortSam being the exception up to 12 Gb of heap space. Effects on GatherVcfs are negligible.

**Fig. 3:**
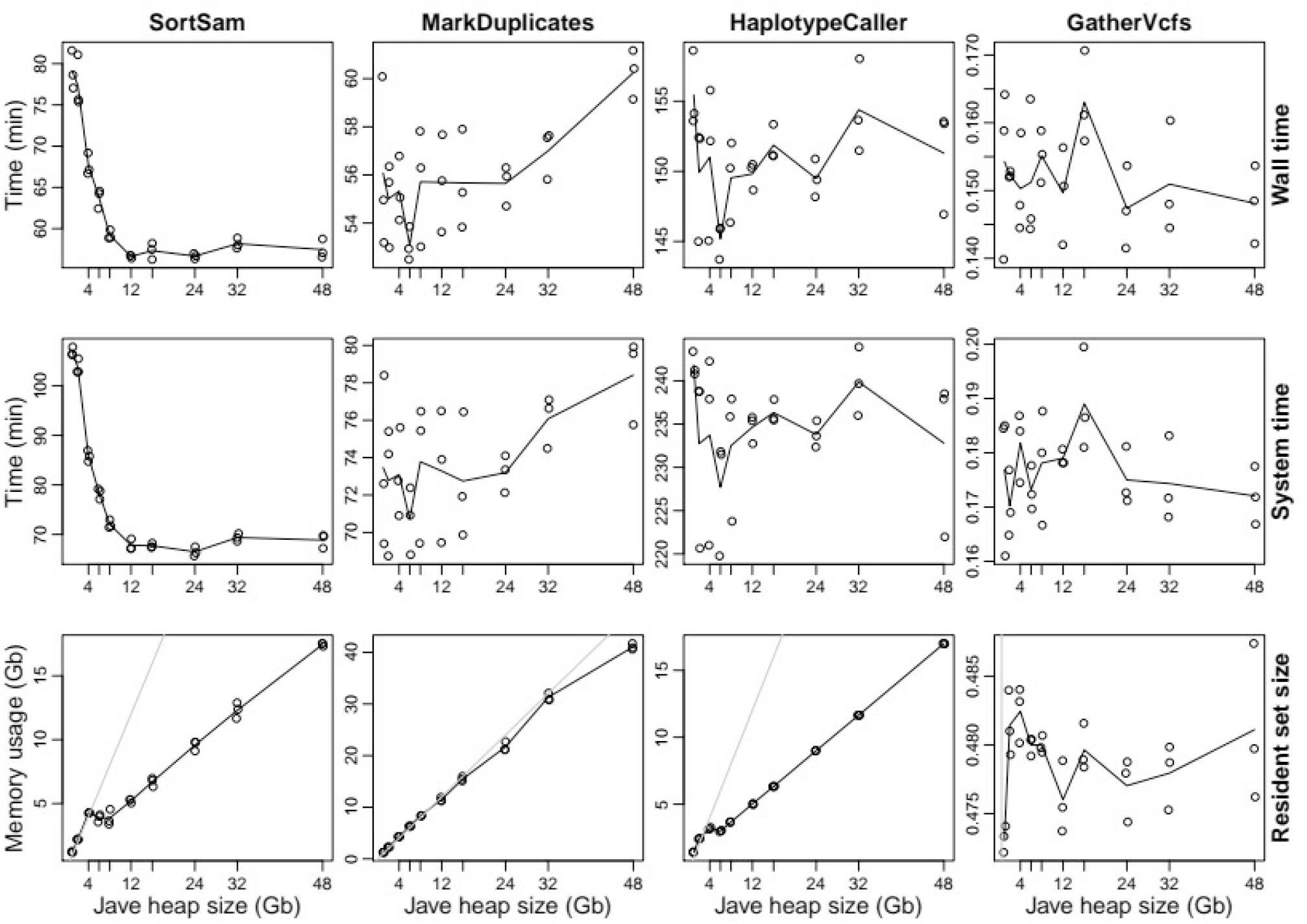
Resource usage of different GATK applications depending on the specified Java heap size. Besides the number of Java garbage collection threads, the provided heap size has a considerable effect on the performance of some GATK applications. Again the four tools SortSam, MarkDuplicates, HaplotypeCaller and GatherVcfs (see columns) (GATK version 4.1.9) were assessed for their respective resource usage concerning wall and system time as well as memory usage (see rows). Triplicated measurements for each of ten different values for the Java heap size (1, 2, 4, 6, 8, 12, 16, 24, 32 and 48 Gb) were recorded and resulting mean values plotted in lines. The gray line in the plots of resident set size indicate parity between maximum allowed heap size and actual memory usage. All measurements were recorded with two garbage collection threads. The different scales of the ordinates of each plot have to be considered, as they vary greatly between the individual plots.

It can be concluded that the heap size exerts drastic impact on memory consumption and in case of SortSam also on CPU usage. To minimize memory usage in the Workflow SortSam was allowed to allocate up to 10 Gb heap space while the other applications were limited to 2 Gb.

### Optimization of variant calling on the workflow level

Previously optimizations were performed on the level of single applications. In this section we want to present how the entire variant calling workflow can benefit from those optimizations. Furthermore we are introducing additional optimizations on the level of parallelized data processing.

CPU usage is only one aspect of hardware utilization. Memory usage can be another limiting factor and shortage thereof can even result in out of memory related application termination. Therefore both of these aspects were monitored during the entire runtime of the workflow [Fig. 4]. For those measurements six fastq files with chicken whole genome re-sequencing data were used. We chose chicken as it’s a vertebrate with a moderately sized genome with approximately 1,1 billion bases.

**Fig. 4:**
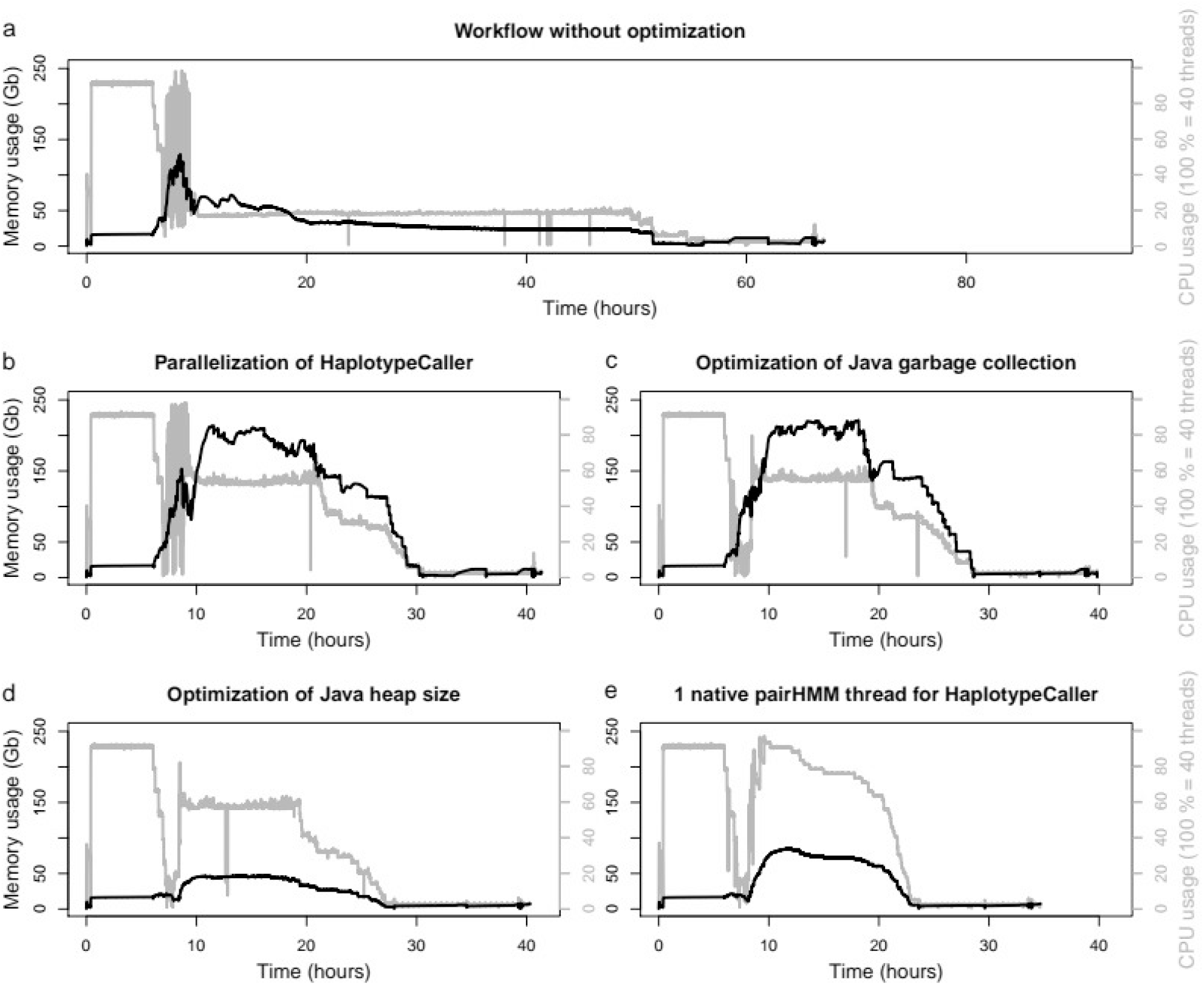
Resource usage of the basic workflow with varying degrees of optimization. **a** CPU and memory utilization of the entire workflow, when using a single interval for the HaplotypeCaller and without any Java optimization (total runtime: 67.1 h)). **b** When the genome is split into six separate intervals for the HaplotypeCaller analysis, but without Java optimization (41.4 h). **c** With optimized Java garbage collection for each GATK application (39.8 h). **d** With optimized Java settings (garbage collection and heap size) for the GATK applications and four default threads for the native pairHMM algorithm of the HaplotypeCaller (40.3 h). **e** When all optimizations are applied to the workflow, including six parallel intervals for the variant calling by the HaplotypeCaller, a single hmmThread for each HaplotypeCaller and all Java optimizations (garbage collection and heap size) (34.7 h).

This shortens the time for workflow benchmarking, while still allowing to estimate evaluation times for larger genomes.

Benchmarking of the workflow was performed for various degrees of GATK optimization. At first (Fig. 4a) a baseline measurement without any particular GATK performance optimizations was recorded. Parallelization was only caused by Snakemake scheduling various processes to be executed simultaneously and by bwa mem to use six threads for each individual mapping process. The plateau in CPU utilization (gray line) within approx. the first six hours is due to mapping by bwa mem. After this phase the workflow is primarily based upon the GATK. Within this phase continuously higher degrees of optimization were implemented.

Firstly parallelization was applied to the HaplotypeCaller (Fig. 4b). This was achieved by splitting the reference genome into six discrete intervals, thereby artificially shortening the reference genome and allowing for more HaplotypeCaller processes to be executed in parallel. In doing so CPU and memory usage are rising during the HaplotypeCaller phase (approx. between 10 and 30 hours), while the overall runtime is shortend by more than an entire day. From this point resource utilization was further reduced through the previously optimized JVM settings.

Applying optimized GC thread values were able to reduce the first spike in memory utilization and caused an reduced and more even CPU usage between 6 and 10 hours of the workflow execution (Fig. 4c). This is mostly related to more efficient resource utilization by SortSam and MarkDuplicates. In the next step usage of the Java heap space was optimized (Fig. 4d). This allowed for a drastic reduction in memory utilization during the phase of parallel execution of SortSam, MarkDuplicates, HaplotypeCaller and GatherVcfs (approx. 6 to 28 h). Before those optimizations were applied, memory utilization maxed out between 230 and 240 Gb and could be lowered to reach a plateau at just about 50 Gb.

The above achieved gains in resource usage paved the way for an additional optimization (Fig. 4e). It was observed, that the HaplotypeCaller, with its default usage of four native pairHMM threads, shows an average CPU utilization of 140 % (additional data is shown in the documentation of OVarFlow, section “Benchmarking & Optimizations » Maximizing CPU utilization”), meaning that one core is fully used and a second core is only used to 40 %. The Snakemake workflow management system on the other hand is only able to calculate and schedule entire threads. This means it can be configured to either use a single or two CPU threads. Therefore CPU usage of the HaplotypeCaller with different numbers of native pairHMM threads was assessed (see the documentation of OVarFlow). This showed that using one native pairHMM thread results in slightly longer runtimes but also only a single thread is utilized. With this setting the Snakemake scheduler only needed to reserve a single thread per HaplotypeCaller, effectively doubling the number of parallel running HaplotypeCaller processes. In doing so CPU utilization could be maximized and the runtime for the entire workflow was further reduced by an additional 5 hours.

In the end all of those optimizations were incorporated into the variant calling workflow, but also were made configurable. The settings can be adjusted through the configuration file config.yml or configBQSR.yml, respectively. This allows for a high degree of customization if needed, while reasonable defaults were set within the workflow itself. As the best performance for an individual HaplotypeCaller is achieved at four native pairHMM threads, this was used as default. This might give a better runtime if unlimited hardware resource are available, e.g. in cluster usage. In this case Snakemake will automatically schedule two threads per HaplotypeCaller process. Still it can be configured to just use a single native pairHMM thread, in which case Snakemake will only reserve a single thread.

## Discussion

The GATK is among the most popular variant calling frameworks [45, 46, 15]. Its Best Practices are commonly referred to as data evaluation procedure, when writing method sections [17, 20]. This often ignores the fact, that the GATK Best Practices are very specifically tailored for human sequencing data [23]. A need for broadly applicable GATK based variant calling workflows is thereby highlighted. Also the Best Practices are continuously evolving, making it hard to reproduce what they recommended some years ago. This is especially problematic with complex data evaluation procedures like variant calling, that involve more than a dozen computational steps and tools (Fig. 1), thereby compromising reproducibility. Badly reproducible methods can also result in a loss of time, due to the requirement to partly reinvent a method or procedure [26]. We therefore created a variant calling workflow, that is oriented towards the GATK Best Practices for germline short variant discovery [47], but is more broadly applicable for model as well as non-model diploid organisms.

The GATK is a very complex framework with hundreds of different tools, were the right tools for variant calling have to be identified first. Additional complexity is introduced through the JVM, that possesses hundreds of configuration options. One major pitfall is, that the JVM does not always scale positively with the size of given hardware resources (see Tab. 2). The JVM tends to allocate more GC threads and larger heap size on a larger hardware base. This in turn can even have negative performance impact on several GATK applications. To many GC threads were slowing down SortSam and MarkDuplicates (Fig. 2) while to large heap spaces consumes more memory than necessary (Fig. 3). In the worst case this can even result in out of memory errors, leading to program termination. The negative impact of too many GC threads for MarkDuplicates is in concordance with previous findings, were various performance optimizations for the GATK were assessed [42]. In this study the authors focused only on the execution time (wall time) of single applications. But this is only one point to consider. System times are equally important, as this better reflects CPU utilization on multithreaded systems. This can for instance be seen with SortSam (Fig. 2), where wall times remain identical no matter the number of GC threads, but system times are increased by approx. 50 % between 1 and 20 GC threads, thereby diminishing the potential for parallel data evaluation. Also memory consumption adds its part to the final bill. Optimization of heap usage reduced memory consumption on the workflow level to less than a quarter (Fig. 3c and 3d), without negative impacts on the total runtime. For one thing this allows to achieve more with given hardware resources, for another thing it might also save costs in cloud environments that consider memory usage in the bill. Here we not only show those performance optimizations but also implemented them in our final workflow, which has not been done in any general purpose GATK 4 based variant calling workflow we are aware of. Besides our improvements to JVM settings, additional system level optimizations were previously assessed for multiple GATK 3.x versions [48]. Applying most of those optimizations require administrator rights, which is often not feasible in multi-user environments. Here we focused on optimizations that can be applied by non-administrator users. Still for large facilities, performing a lot of variant calling, this might be an opportunity to further accelerate variant calling.

Having a clearly defined data evaluation pipeline is only one aspect of scientific reproducibility. The most reproducible workflow is of no help, if the underlying software cannot be installed. Mangul *et al*. performed a systematic analysis of software installability, in which 51 % of the investigated tools were claimed “easy to install”, but still 28 % failed to be installed at all [28]. To circumvent installation obstacles and to promote the long-term usefulness of our workflow, we followed the generally guidelines laid out by Manguel *et al*. Firstly our software is hosted publicly by GitLab and the accompanying documentation is available through Read the Docs. Secondly installation relies on the well established Conda environment and package manager. Thirdly Conda also takes care about dependencies of all used tools. Furthermore the last two points are also guaranteed by providing container images, even further reducing any required efforts on the side of the end user. Additionally the documentation not only includes an example dataset, but a detailed tutorial on how to evaluate this dataset. Besides a quick start guide a detailed description of every software component used is included. The software can be run without any root privileges. The docker container might even be run on a Windows system. However, we strongly recommend performing variant calling on Linux based high performance computing infrastructure.

Finally our variant calling workflow is entirely based upon open source software. This is a vital point for the unlimited use and availability of our workflow. License changes between GATK 3 and 4 made this possible. With the previous license of GATK 3, distribution of an prepackaged workflow in a a container environment would not have been possible. This highlights the value of the open source software and its related licensing schemes, to the scientific community.

## Conclusion

Variant calling has become an established method in genomics research. Its importance can only be expected to rise due to continuously declining sequencing costs. The GATK, developed at the Broad Institute, is a major player in the field and its Best Practices are commonly cited in related publications, despite being focused on human sequencing data. This shows the high demand for an easy accessible and broadly applicable GATK based variant calling workflow. Our workflow perfectly fits into this niche, offering automatization, reproducibility and documentation of the variant calling process. We are confident that our developed workflow will help filling this gap and further lower the threshold for variant calling, also for smaller institutions. Furthermore we hope that our presented optimizations not only help to increase throughput, but also to utilize given hardware resources more efficiently.

## Methods

### Utilized software versions

The following software versions were used within the variant calling workflow: FastQC v0.11.9, bwa 0.7.17-r1188 [36], samtools 1.11 [49], GATK 4.1.9 [16] and SnpEff 5.0 [34] (in the order of usage). Of those FastQC, GATK and SnpEff rely on the JVM for program execution. Here the OpenJDK version 1.8.0_152 was utilized. Snakemake [31] version 5.26.1 acted as a workflow management engine. Software installation was performed via Conda 4.9.2. Default JVM resource usage on various machines was determined using the commands `java -XX:+PrintFlagsFinal | grep ParallelGCThreads` and `java -XshowSettings:vm`.

### Data analysis for dwarf chicken variant

Whole genome sequencing data were obtained from the ENA, for one dwarf chicken (ERR2505843) and 25 White Leghorns (ERR3525251-8, ERR4326765-74, SRR2131198-9, SRR2131201, SRR5079491-3, SRR5079496). The variant calling workflow was executed using chicken genome build Gallus_gallus-5.0. To reduce the total runtime, all contigs shorter than 20,000 bp were excluded. Variants on chromosome 1, within the candidate region 52–56 Mb, were extracted by the bcftools (1.6) view command. Filtering of potentially causal variants was performed using a custom Python 3 script, filtering for homozygous variants exclusive to the dwarf chicken. Moderate and high impact variants, as categorized by SnipEff, were selected using the Unix command `grep ‘\(MODERATE\|HIGH\)’`.

### Benchmarking of individual GATK applications

To reduce application runtime, sequencing data from *Gallus gallus* were employed to benchmark the performance of single GATK applications. The current representative genome GRCg6a (RefSeq assembly accession: GCF_000002315.6) was obtained from the RefSeq. Fastq files were obtained from the ENA, run accession SRR3041137, offering 2 x 125 bp Illumina sequencing data (HiSeq 2500). For this dataset an average coverage of 34 with standard deviation of 44 was determined (see rule `calculate_average_coverage` of the workflow for details). SortSam, MarkDuplicates and GatherVcfs processed the entire dataset, while HaplotypeCaller was restricted to the contig `NC_006093.5` (36,374,701 bp) to reduce application runtime. To benchmark various Java GC thread numbers the heap size was fixed to 12 Gb. For benchmarking the effect of various Java heap sizes, two GC threads were specified. Resource usage of each application was monitored by GNU time version 1.8 (version 1.7 reports incorrect results), written to a log file and visualized using R (3.4.4). Resource measurements were recorded employing page caching by copying all accessed data to `/dev/null` before actual program execution. Especially for a short running application like GatherVcfs runtimes will be noticeably longer without page caching.

### Benchmarking of the entire workflow

Again reference genome and annotation GRCg6a of *Gallus gallus* were used. Paired end Illumina sequencing data (2 x 125 bp) were downloaded from the ENA, project PRJEB12944, run accessions ERR1303580, ERR1303581, ERR1303584, ERR1303585, ERR1303586 and ERR1303587. Average coverage of those files was between 24 and 28. Five intervals were specified in the `config.yaml` file for parallelization of the HaplotypeCaller (`GatkHCintervals`). Thereby the reference genome was actually split into six discrete intervals. This is intended behavior of the splitting algorithm, which only splits the reference genome between individual contigs, while trying to limit the maximum size of the created intervals (see `createIntervalLists.py` for implementation). To measure resource usage of the entire workflow the commands `mpstat 30` and `sar -r ALL 30` from the sysstat application suit (12.2.0) were employed. Recorded measurements were plotted using R (3.4.4).

## Availability and requirements

|  |  |
| --- | --- |
| Project name: | OVarFlow |
| Project home page: |  |
| - GitLab: | https://gitlab.com/computational-biology/ovarflow |
| - Read the Docs: | https://ovarflow.readthedocs.io/en/latest/ |
| - Docker hub: | https://hub.docker.com/r/ovarflow/release/tags |
| - Zenodo (Singularity): | https://zenodo.org/record/4746639 |
| Operating system(s): | tested and working on Linux<br>container images might also work on macOS and Windows |
| Programming language: | Python & Snakemake; Bash |
| Other requirements: | Conda/Bioconda or Docker/Singularity |
| License: | Code: GNU General Public License version 3 (GPLv3) |
|  | Documentation: Creative Commons license CC-BY SA 3.0 |
| Any restrictions to use by non-academics: | none |

### List of abbreviations

Adw: autosomal dwarfism
AVX: Advanced Vector Extensions
BQSR: base quality score recalibration
bwa: Burrows-Wheeler Aligner
CPU: central processing unit
csv: colon separated values
ENA: European Nucleotide Archive
GATK: Genome Analysis Toolkit
Gb: gigabytes
GC: garbage collection
GWAS: genome-wide association study
JVM: Java Virtual Machine
NGS: next generation sequencing
RSS: resident set size
SGS: second generation sequencing
SNP: single nucleotide polymorphism
VCF: variant call format
WES: whole exome sequencing
WGS: whole genome sequencing

## Declarations

## Acknowledgments

We want to thank the Institute for Bioinformatics and Systems Biology at the Justus Liebig University Giessen for providing computational resources and support.

## Ethics approval and consent to participate

Not applicable.

## Consent for publication

Not applicable.

## Availability of data and materials

See above for availability of code, documentation and containerized software images.

## Authors’ contributions

Jochen Bathke devised the workflow, performed the benchmarking, wrote the code and documentation. Jochen Bathke and Gesine Lühken conceived the project and wrote the manuscript.

